# SECRET AGENT O-GlcNAcylates hundreds of proteins involved in diverse cellular processes in Arabidopsis

**DOI:** 10.1101/2023.09.16.558074

**Authors:** Ruben Shrestha, Sumudu Karunadasa, Tara Grismer, Andres Reyes, Shou-Ling Xu

## Abstract

O-GlcNAcylation is a critical post-translational modification of proteins observed in both plants and animals and plays a key role in growth and development. While considerable knowledge exists about over 3000 substrates in animals, our understanding of this modification in plants remains limited. Unlike animals, plants possess two putative homologs: SECRET AGENT (SEC) and SPINDLY (SPY), with SPY also exhibiting O-fucosylation activity. To investigate SEC’s role as a major O-GlcNAc transferase in plants, we utilized LWAC enrichment and SILIA labeling, quantifying at both MS1 and MS2 levels. Our findings reveal a significant reduction in O-GlcNAc levels in the *sec* mutant, indicating SEC’s critical role in mediating O-GlcNAcylation. Through a comprehensive approach, combining HCD and EThcD fragmentation with substantial fractionations, we expanded our GlcNAc profiling, identifying 436 O-GlcNAc targets, including 227 new targets. The targets span diverse cellular processes, suggesting broad regulatory functions of O-GlcNAcylation. The expanded targets also enabled exploration of crosstalk between O-GlcNAcylation and O-fucosylation. We also examined EThcD fragmentation for site assignment. This report advances our understanding of O-GlcNAcylation in plants, facilitating further research in this field.

## Introduction

Protein O-GlcNAcylation is a unique post-translational modification (PTM) involving the addition of a single monosaccharide, β-N-acetylglucosamine (GlcNAc), to the hydroxyl group of serine or threonine residues on nuclear, cytoplasmic, or mitochondrial proteins. This modification, discovered in both animals and plants, has been shown to play an essential role in growth and development (1, 2). O-GlcNAcylation has been demonstrated to participate in a diverse range of cellular processes, including signal transduction, transcriptional regulation, stress response, cellular differentiation, and protein degradation. Dysregulation of this PTM has been linked to various diseases, including Alzheimer’s disease, cardiovascular disease, diabetes, and cancer (3).

O-GlcNAcylation is catalyzed by O-GlcNAc transferase (OGT), comprising an N-terminal tetratricopeptide repeat (TPR) domain for protein interactions and a C-terminal catalytic domain. OGT acts as a nutrient sensor (4), responding to hexosamine biosynthetic pathway flux and UDP-GlcNAc levels (5). Complete OGT knockout is embryonically lethal in various animal models, while knockdown or conditional knockout reveals diverse phenotypic functions. Human OGT (hOGT) exhibits catalytic activity in vivo and in vitro, being the sole enzyme responsible for these PTMs.

In contrast to hOGT, plants have two closely related enzymes, SECRET AGENT (SEC) and SPINDLY (SPY). Knocking out of SPY leads to mild effects, including a spindly shoot, reduced GA sensitivity, male sterility, early flowering and altered phyllotaxy (6). *sec* mutants are minimally affected, except for some impact on virus infection efficiency (7) and early flowering (8). Double mutants of SEC and SPY are lethal (9, 10). In vitro, SEC has shown activity using *E.Coli* (10), whereas SPY lacks such activity. However, SPY exhibits O-fucosylation activity both in vitro and in vivo (11). Extensive quantification studies comparing wild-type and *spy* mutants, after enriching O-fucosylated peptides, highlight SPY as the primary enzyme responsible for O-fucosylation activity (12, 13).

Recently, a discovery in wheat revealed an atypical TaOGT(TaOGT1) with no structural similarity to SEC and SPY enzyme (14). TaOGT1 was found to O-GlcNAcylate TaGRP2. In Arabidopsis, approximately 34 unannotated or uncharacterized proteins share similarity with TaOGT1, leading to questions about whether SEC is the primary contributor to O-GlcNAcylation in plants.

Comparatively, although over 3000 and 1430 O-GlcNAc targets have been identified in humans and mice, respectively (15, 16), the database for O-GlcNAc targets in plants remains limited (17–20). Additionally, given the overlapping or even opposite functions of O-GlcNAcylation and O-fucosylation, identifying more common targets would help in understanding their roles. Previously, we identified 262 O-GlcNAc targets (20)and 467 O-fucosylated targets in Arabidopsis (12).

In this study, we conducted genetic, quantitative proteomic, and large-scale proteomic experiments to investigate O-GlcNAcylation. We confirmed that SEC and SPY are essential for embryo cell viability. Through quantitative proteomics, we demonstrated that SEC is a major contributor to O-GlcNAcylation in plants. Moreover, we significantly expanded the O-GlcNAc target identifications, discovering a total of 436 O-GlcNAc targets, including 227 new targets. Additionally, we explored the crosstalk between O-GlcNAcylation and O-fucosylation. Finally, we delved into the complexity of site assignment using EThcD for O-GlcNAc studies.

## RESULTS

### Embryo lethality of knocking down both *sec* and *spy*

Previous genetic studies have shown that SEC and SPY have redundant functions during gametogenesis and embryogenesis, despite *sec* single mutant not displaying any severe phenotype (10). This was demonstrated by constructing double mutants using the *spy-3* (point mutation in the catalytic domain, AT3G11540) and *sec-1* or *sec-2* (AT3G04240) allele and not identifying the double mutant alleles in F2 generation. To further investigate this functional redundancy, we attempted to construct a *sec spy* double mutant by crossing *spy3* with *sec-5* (8). This *sec-5* allele carries an insertion in the second exon of the SEC genome, resulting in a significant decrease in SEC transcript levels and is considered the strongest allele. As SEC and SPY are genetically linked due to being close on the same chromosome, we screened 520 seeds collected from *sec5*(-/-) *spy3*(+/-) plants for germination on paclobutrazol (PAC)-containing plates and expected to get ¼ of seeds that would be double mutant,which would be PAC resistant. We failed to recover any double mutants (data not shown). We also grew these seeds on ½ MS plates and genotyped plants but failed to identify any double mutants (data not shown). We next examined the siliques and observed that while *spy-3* and *sec-5* individually exhibited similar embryos to the wild-type (WT), siliques from *sec-5*(-/-) *spy3*(+/-) produced 17-20% of embryos that turned brown and shrank to a smaller size (Fig. 1). Additionally, we observed shrunken seeds in the mature stage (Fig. 1) that failed to germinate (data not shown). These findings confirm that SEC and SPY have overlapping functions in embryo development, and highlight the importance of O-Glycosylation, including both O-GlcNAcylation and O-fucosylation, for embryo cell viability.

**Fig. 1.**
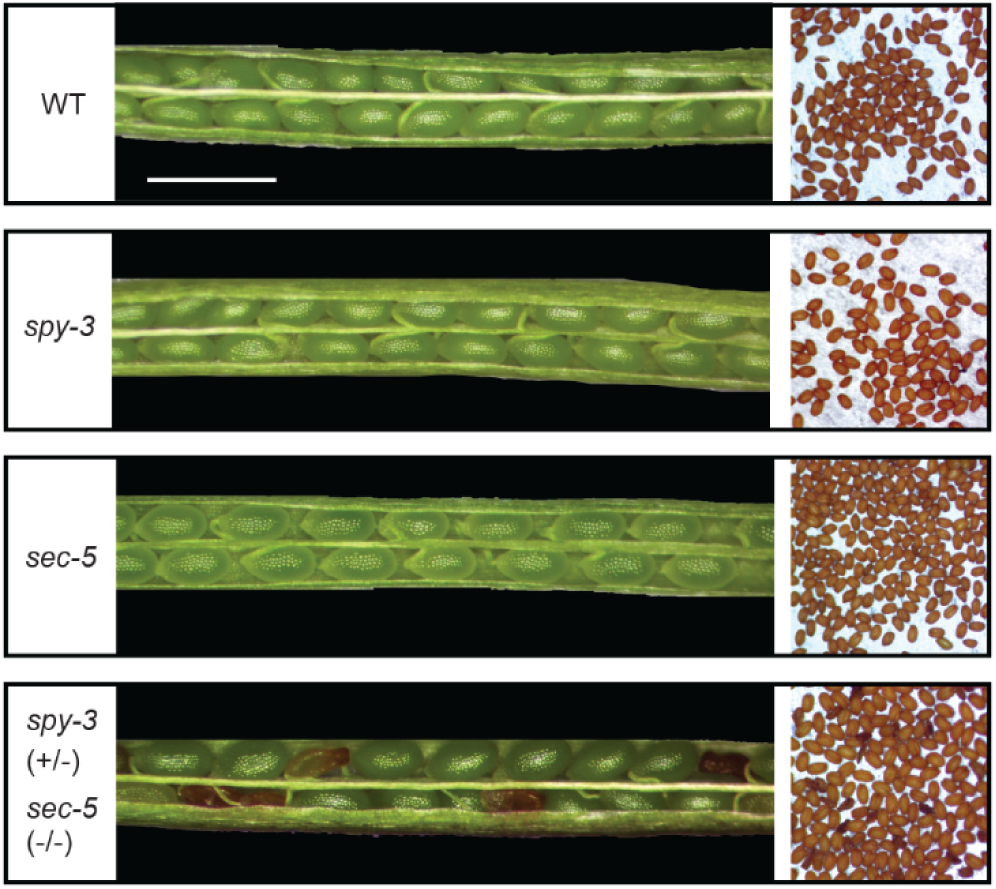
Essential functions of SEC and SPY in embryo development. Representative siliques from wild type (WT), *spy-3, sec-5, sec-5(-/-) spy-3(+/-)* plants are shown on the left, and seed set for each genotype is shown on the right. Siliques from *sec-5*(-/-) *spy3*(+/-) plants produce approximately 17-20% of embryos exhibiting browning and reduced size, which persist throughput maturity. Scale bar =1mm.

### Quantifying O-GlcNAc abundance levels between wild type vs *sec-5* mutant

Next, we determined if SEC is the major contributor to O-GlcNAcylation. We conducted a comparison of O-GlcNAc-modified peptide levels between WT and *sec* mutant samples. To enrich for O-GlcNAc-modified peptides, we utilized the lectin-weak affinity chromatography (LWAC) pipeline as described in our previous study (20). We then quantified the levels of O-GlcNAc between WT and *sec-5* mutant using a workflow involving stable isotope labeling in Arabidopsis (SILIA) and quantified using both data-dependent acquisition (DDA) and targeted quantification via Parallel Reaction Monitoring (PRM) (21, 22) (Fig.2A).

**Fig. 2.**
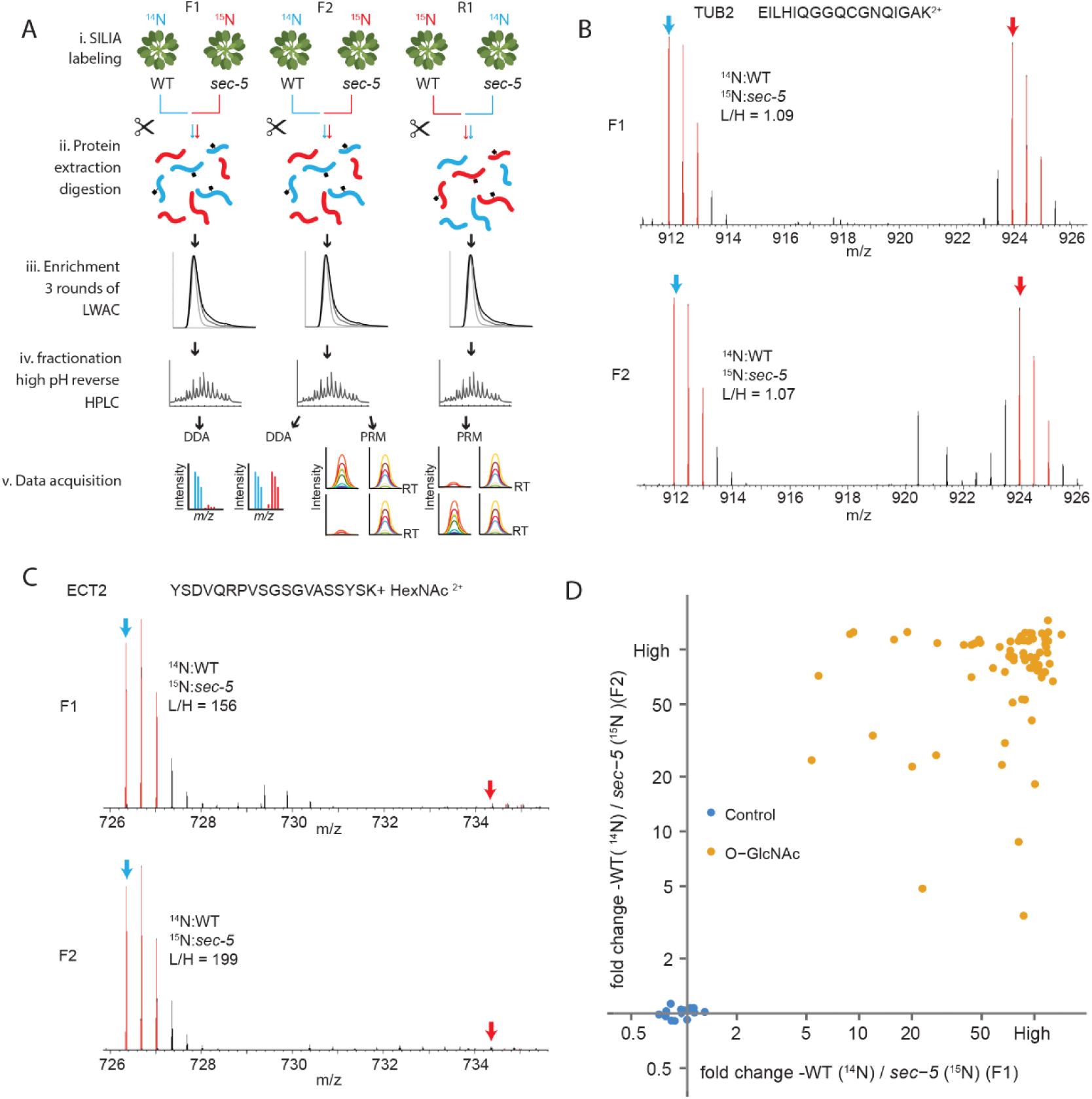
Quantitative workflow to determine SEC’s role in O-GlcNAcylation in Arabidopsis. A. Workflow for stable isotope-labeling, LWAC-enrichment, high-pH reverse HPLC fractionations, and mass spectrometry (MS) analysis. Total proteins from ^14^N- and ^15^N-labeled wild-type (WT) plants and *sec-5* mutants were mixed, extracted, and digested with trypsin. For each mixture, O-GlcNAc-modified peptides were captured through 3 rounds of LWAC enrichment and separated by high pH reverse HPLC fractionations. LC-MS/MS analysis was performed using data-dependent acquisition (DDA) and parallel reaction monitoring (PRM). B-C. Selected MS1 spectra illustrating the relative abundance of non-modified peptides and O-GlcNAc-modified peptides in WT compared to *sec-5* mutant (F1 and F2 replicates). Spectra represent peptides from TUBULIN BETA CHAIN2 (TUB2, AT5G62690) (B) and EVOLUTIONARILY CONSERVED C-TERMINAL REGION 2 (ECT2) (C). D. Scatterplot showing ratios of O-GlcNAc-modified peptides from modified proteins and control peptides from housekeeping proteins. Quantification using SILIA-MS1 was performed in both F1 and F2 replicates comparing WT vs *sec-5* mutants.

Briefly, we grew WT and *sec-5* mutant seedlings on media containing the ^14^N and ^15^N stable isotopes (F1 & F2 biological replicates, ^14^N WT and ^15^N *sec-5*) and harvested the plants on day 14. In the third biological replicates (R1), we reversed the isotope labeling (R1, ^15^N WT and ^14^N *sec-5*) (Fig. 2A). After protein extraction from mixed tissues and tryptic digestion, each peptide sample was enriched for GlcNAc-modified peptides using three rounds of LWAC and subjected to high pH HPLC fractionation. All peptide samples were analyzed by liquid chromatography coupled with high-resolution and high-accuracy orbitrap Q-Exactive HF or Orbitrap Eclipse (LC-MS/MS). The F1 and F2 replicates were analyzed using DDA and quantified at the MS1 level, while F2 and R1 were quantified using PRM at the MS2 level (Fig. 2A).

The SILIA DDA data from F1 and F2 give high-confidence quantification data. The equal mixture of ^14^N-labeled WT and ^15^N-labeled *sec-5* was exemplified in TUBULIN2 quantification (F1, L/H =1.09; F2, L/H=1.07) (Fig. 2B). However, the O-GlcNAc-modified peptide from EVOLUTIONARILY CONSERVED C-TERMINAL REGION2 (ECT2) (Fig. 2C) showed a significant reduction in the *sec-5* mutant compared to WT (F1, L/H=156; F2, L/H=199). In total, we quantified 72 O-GlcNAc peptides from 57 proteins detected in both replicates (Fig. 2D, supplementary data 1). Apart from TUB2, 12 other housekeeping proteins also verified the equal mixture between wild-type and *sec-5* (Fig. 2D). In contrast, the scatter plot demonstrated that all quantified O-GlcNAc modified peptides detected in both replicates consistently exhibited a significant reduction in the *sec-5* mutant (Fig. 2D).

Next, we performed targeted quantification using PRM in the F2 and R1 replicates. PRM overcomes the low identification issues in ^15^N labeled samples as well as missing values in replicates (22). To select transitions, we chose y-type ion fragments due to their higher ion abundance compared to b-type ion fragments (23). We quantified 10 peptides with 4-5 transitions, while all the rest of peptides used ≥ 6 transitions to quantify the abundance. With these stringent criteria, we obtained high confidence in quantification. Consistent with MS1 quantification, PRM showed the control TUBULIN2 peptide having equal abundance between WT and *sec-5*. In contrast, O-GlcNAc peptide PRM shows a striking reduction in *sec-5* mutant, as exemplified by a modified peptide from the AT-HOOK MOTIF NUCLEAR LOCALIZED PROTEIN 1 (AHL) (Fig. 3B). In total, we quantified 172 O-linked peptides from 109 proteins using skyline quantification (22, 24). Targeted quantification shows consistent reduction in the *sec-5* mutant, with only 2 O-GlcNAc modified peptides showing about 3-fold median difference, while 170 show a significant difference (>5-fold median) (Fig 3C, supplementary data 1) in which 163 peptides show over 25-fold difference. In contrast, the N-linked peptides did not show consistent quantification difference across three replicates (supplementary Fig. 1).

**Fig. 3:**
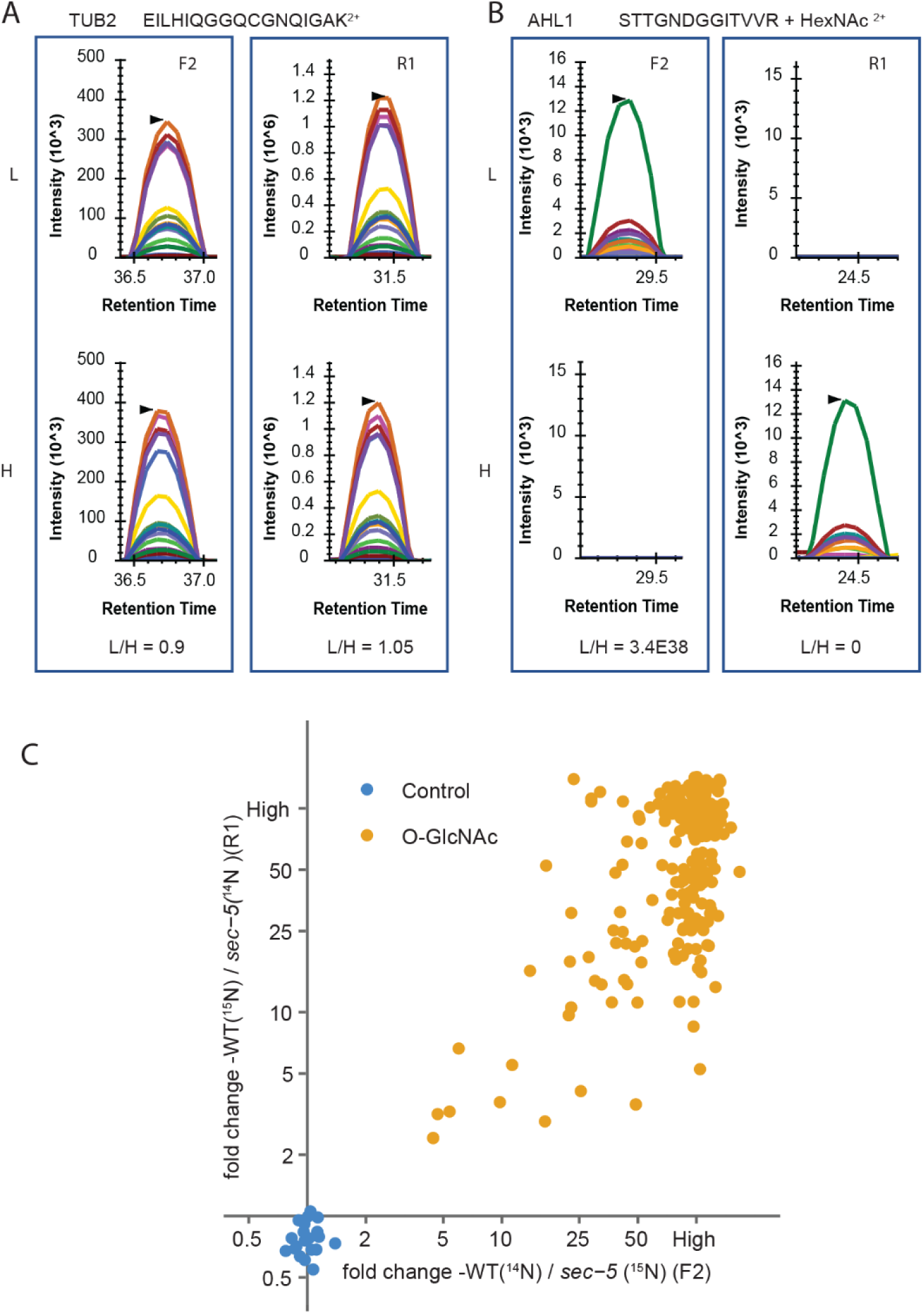
Targeted quantification reveals reduced O-GlcNAcylation in *sec-5* mutant. A. PRM data demonstrates equal abundance of TUB2 peptides between WT and *sec-5* mutants in F2 and R1 replicates. B. O-GlcNAc modified peptides from AT-HOOK MOTIF NUCLEAR-LOCALIZED PROTEIN 1 (AHL1) protein show a significant reduction in *sec-5* mutant. Selected transitions were extracted for the ^14^N labeled (L) and ^15^N-labeled (H) peptides. Area under the curve was used for ratio calculation. C. Scatterplot displays ratios of O-GlcNAc-modified peptides and control peptides using SILIA-PRM. Quantification was performed in both F1 and R1 replicates, comparing WT vs *sec-5* mutants.

Taken together, the quantitative results from these replicates on 184 peptides across 115 proteins using DDA and PRM suggest that SEC is a major contributor to O-GlcNAcylation in Arabidopsis.

### Large-scale O-GlcNAc peptide identifications

The O-GlcNAcome database has comprehensively cataloged over 3000 and 1430 O-GlcNAc targets identified in humans and mice (15, 16). However, the database for O-GlcNAc targets in plants is still relatively limited. We previously reported 262 O-GlcNAc-modified proteins found in Arabidopsis inflorescence tissue. To further expand our knowledge of the O-GlcNAcome in plants, we performed a series of comprehensive mass spectrometry runs, totaling 107 data-dependent acquisitions (DDA) measurements.

In our measurement, we employed two acquisition approaches: the higher-energy collision dissociation (HCD) method on Q Exactive HF and the HCD/204 product-dependent electron-transfer high-energy collision dissociation (EThcD) analysis on Tribrid Eclipse. HCD is sensitive and effective in confidently identifying O-GlcNAc peptides, thanks to its short cycle time and efficient backbone fragmentation. However, it cannot assign modification sites due to the neutral loss of the very labile O-GlcNAc moiety from fragment ions. Conversely, ETD enables site assignments by producing c/z backbone fragment ions that retain the O-GlcNAc moiety, facilitating mass shifts for site localization (25). Nonetheless, ETD may lead to EtnoD, reducing the sensitivity of detecting O-GlcNAc-modified peptides (26).

To overcome these challenges, we utilized HCD/204pd-triggered EThcD, which provides a richer spectrum and yields more site assignments (27). Notably, fragments obtained from ETD activation retain the modification in EThcD, while those from HCD activation often do not for O-GlcNAc peptides. We therefore employed two methods, HCD-m or EThcD-m1, and EThcD-m2, for data searches. HCD-m or EThcD-m1 allows both loss and retention of the modification (Serine/Threonine/Asparagine (S/T/N)) on fragments (see methods) for data acquired using HCD and EthcD; HCD-m considers b, y ions while EtHCD-m1 takes into account c, z, b, y ions. Our in-house software Protein Prospector treats them as separate options and reports the higher scoring result. EThcD-m2 is used for EThcD data, and assumes all fragments are modified on S/T/N and considers c, z, b, y ions.

To ensure the high-confidence identification of O-GlcNAc modified proteins, we implemented several criteria for data filtering. These included examining if MS2 contains a 204-oxonium ion peak, or if peptides are identified by HCD and EThcD, or if modified peptides contain different miscleavage. We found several exceptions, such as the 204 ions being out of the scanning range in MS2 spectra for some peptides with a larger mass (over 2000). To address these cases, we manually examined their spectra and observed that the MS2 spectrum often accompanied the precursor-neutral loss of sugar (-203.08).

Our mass spectrometry analysis of LWAC-enriched peptides yielded a total of 1721 O-GlcNAc peptides, corresponding to 1057 unique peptide sequences derived from 436 proteins (Supplementary data 3). When compared to our previous study that utilized LWAC-enriched samples from Arabidopsis inflorescence tissue (20), we discovered 227 new O-GlcNAc-modified proteins (Supplementary data 4). However, 53 proteins were not detected in this study, likely due to their specific expression in flower tissue.

The data obtained from EThcD allowed us to successfully identify 596 O-GlcNAc modification sites in peptides (supplementary data 3). Sequence analysis of O-GlcNAcylation sites did not reveal any consensus sequences, except for a slight preference for proline at -2 or -3 positions N-terminal, a slight preference for valine residues at -1 position to the modification site. Furthermore, serine residues were often observed at positions C-terminal to the modification sites (Fig. 3A). This finding is consistent with previous reports in (20) as well as reports from animal studies (16, 28). These results align with the structure analysis of human OGT, indicating that it primarily binds to the peptide backbone of substrates and shows no clear specificity for the modifications of specific serine or threonine (29).

Given no clear consensus, to our surprise, we discovered O-GlcNAcylation occurring in various members of the same protein family (Supplementary data 4). For instance, we identified 20 modified family members belonging to the CCCH-Type Zinc finger family, 12 members from the AT HOOK MOTIF DNA-binding proteins family, 9 members from the ENTH/VHS family proteins, 9 members from TCP family transcription factor, 8 members from the Evolutionarily conserved C-terminal region proteins, 8 members from the bHLH DNA-BINDING family proteins, 8 members from the homeodomain containing proteins, 6 members from the PUMILIO FAMILY, 5 members from the OCTICOSAPEPTIDE/PHOX/BEM1P FAMILY PROTEIN, 5 members from the AUXIN RESPONSE FACTORs, and 3 members from the GYF domain-containing proteins.

We proceeded to examine the specific regions where these modifications occur within these proteins. Interestingly, despite the absence of the consensus sequences derived from the motif assay, we discovered that similar regions within the same protein family are subjected to modification (Fig. 4B). For example, in the mRNA binding protein family PUMILIO, we observed that PUMILIO 3 (At2G29140), along with its homologs PUMILIO 1 (At2G29200) and PUMILIO 2 (At2g29190), are all modified at the same region, as indicated by the blue-colored modified peptides. Likewise, in the auxin transcription factors, we found that both AUXIN RESPONSE FACTOR 8 (ARF8, At5G37020), and its homolog ARF6 undergo O-GlcNAc modification in their N-terminal regions. Additionally, we observed that similar regions of the heavily modified GYF protein family (AT1G27430, AT1G24300) are also subject to modification (the shared identical peptides are not displayed here) (Fig. 4B). Based on these findings, we hypothesize that there may be adaptor proteins employed by the SEC, similar to the proposed mechanism for hOGT, enabling the modification of different members from the same family (30). Further exploration of the O-GlcNAcome with even higher coverage will provide more opportunities for comparison and deeper insights into the OGT selectivity and specificity.

**Fig. 4:**
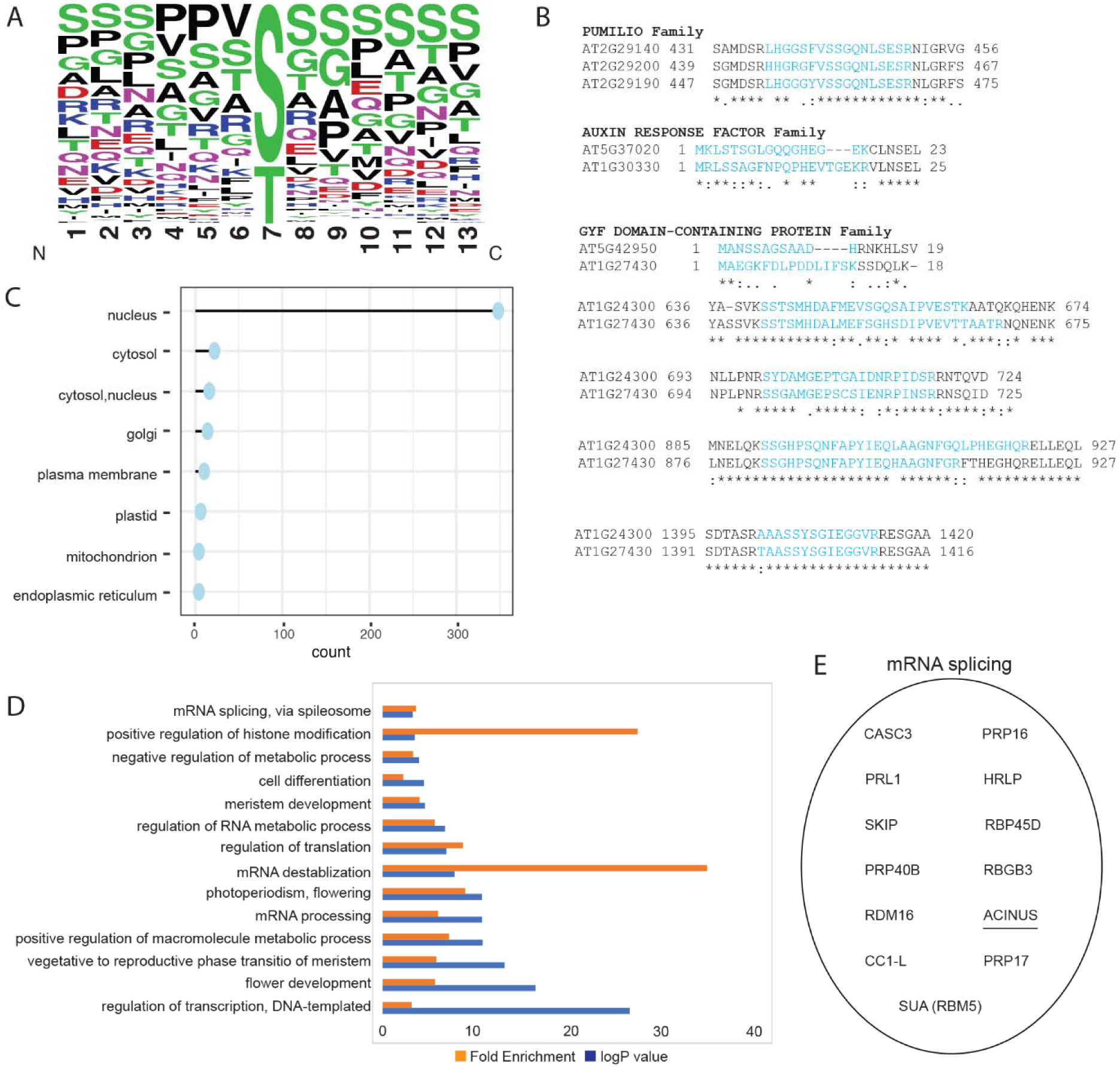
Summary of identified O-GlcNAcylated proteins and peptides. A. Weblogo motif representing the alignment of identified O-GlcNAc modified sites. B. Comparison of selected modified regions within the same modified family. C. SUBA analysis reveals predominant localization of O-GlcNAcylated acceptor-substrate in the nucleus and cytosol. D. Gene Ontology (Go) analysis of in vivo detected O-GlcNAcylated proteins. Yellow indicates fold enrichment, and blue indicates log p-value for biological processes on the y axis. E. Novel mRNA splicing factors are identified. Previously identified splicing factor ACINUS is underscored.

### SUBA and GO analysis of O-GlcNAc substrates

Subcellular localization analysis of the O-GlcNAc proteins using SUBA5.0 (31) revealed that the majority of the O-GlcNAc modified proteins are predicted to be localized in the nucleus (Fig.4C,supplementary data 4). The next most abundant group of O-GlcNAc modified proteins is predicted to be in the cytosol, while only a small number of proteins are predicted to be localized elsewhere. However, we noted that some SUBA predictions may not be accurate. For example, At2g32080, PURIN-RICH ALPHA1, is predicted by SUBA to be localized to Golgi, but TAIR predicts to be active in the nucleus. Considering its O-GlcNAc modification and its homologs’ involvement in transcriptional control and cytoplasmic RNA localization in animals(32), it is more likely to be cytosolic or nuclear.

Gene Ontology analysis revealed a significant enrichment of crucial molecular functions associated with O-GlcNAc modified proteins (Fig.4D). These functions encompassed mRNA splicing, histone modification, cell differentiation, meristem development, translation, mRNA destabilization and processing, as well as transcription. Notably, a substantial portion of the identified O-GlcNAcome was found to be involved in RNA metabolism, histone modification and translation, which mirrors observations in the human O-GlcNAcome, suggesting a conserved regulatory mechanism for O-GlcNAcylation in these processes.

mRNA splicing components are notably enriched in our findings (Fig.4D and 4E). Specifically, in addition to previously reported ACINUS, we identified 12 additional novel splicing factors, including CASC3, PRP16, PRL1, HRLP, SKIP, RBP45D, PRP40B, RBGB3, RDM16, CC1-L, PRP17 and SUA(RBM5). We previously showed that both SEC and SPY can affect splicing in Arabidopsis (33), and similar effects on splicing have also been observed when O-GlcNAc is perturbed in animals (34).

Furthermore, our analysis revealed a significant enrichment pointing to the central role of O-GlcNAc in signaling. For instance, we identified seven proteins harboring the WD40 (including NEDD1, VCS, TWD40-1, KTN80 Subunit 4, FY, PRP17, LUG) and seventeen proteins with kinase domains as prominent targets of O-GlcNAcylation. These WD40 and kinase domains exhibit extensive interaction surfaces and are commonly associated with vital functions in signal transduction pathways.

### Crosstalk between O-GlcNAcylation and O-fucosylation

In our study, we identified a total of 436 O-GlcNAc targets, including 227 newly discovered targets, which complements the previously reported 262 targets and expands the total number of identified O-GlcNAc targets to 489 (Fig. 5A). Given both the known shared and distinct functions of SEC and SPY (35), we compared the O-GlcNAc and O-fucosylated datasets and were able to expand the shared targets from 128 (12) to 184, suggesting a strong potential crosstalk between these two PTMs, with over ⅓ of the targets overlapping between these two datasets.

**Fig 5.**
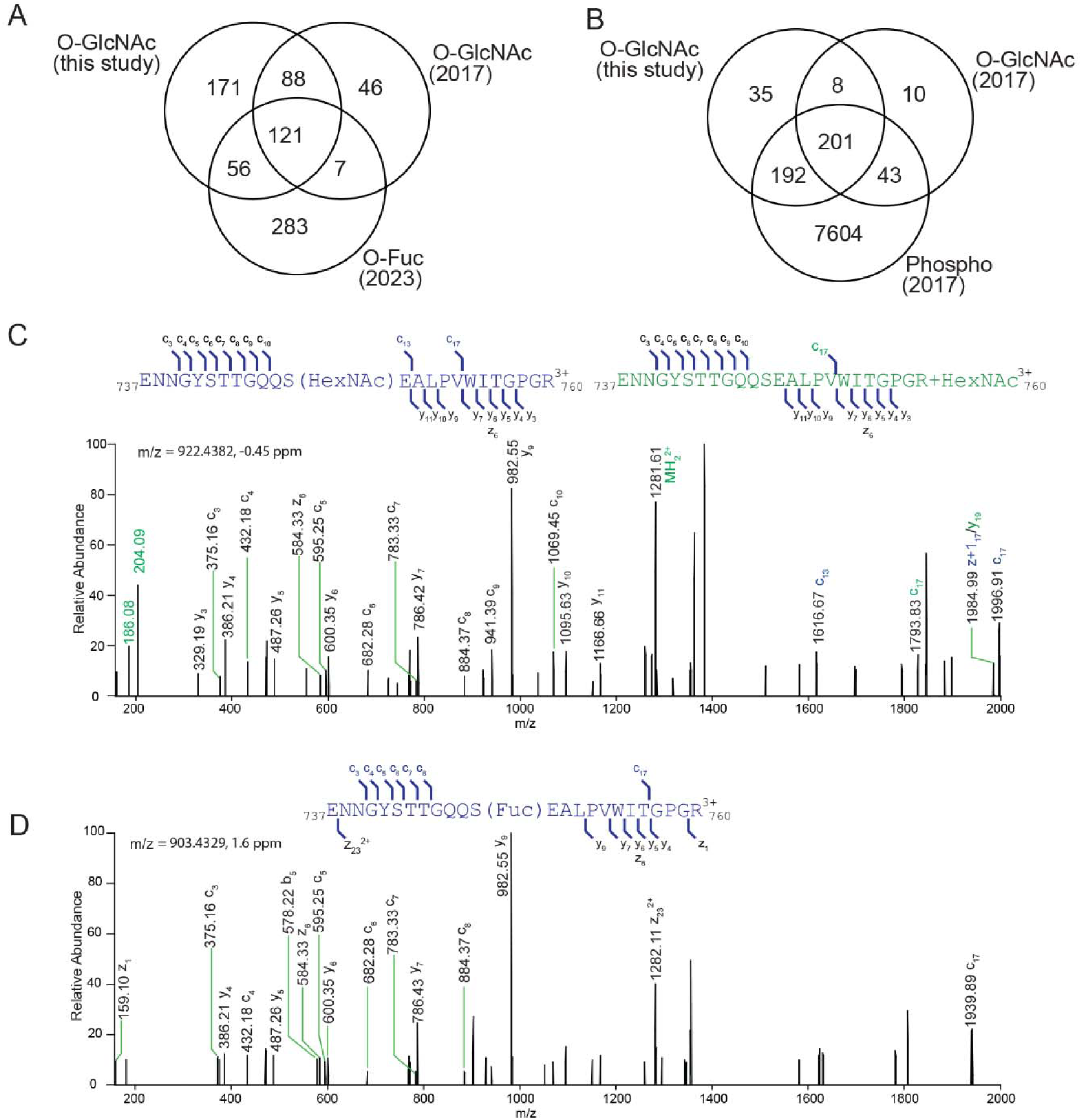
Crosstalk between O-GlcNAcylation, O-fucosylation and O-phosphorylation. A. A Venn diagram illustrates the intersection of O-GlcNAcylated proteins identified in this study, previously reported O-GlcNAcylated proteins, and O-fucosylated proteins. B. A Venn diagram demonstrates the overlap between O-GlcNAcylated proteins and O-phosphorylated proteins. C-D. EtHCD spectra of m/z 922.4382 and m/z 903.4329 identify a peptide from KINASE-RELATED PROTEIN OF UNKNOWN FUNCTION (AT3G07660) being modified by O-GlcNAcylation or O-fucosylation at the same Serine 748 site. In (C), EtHCD spectra display fragments that contain modifications (highlight in blue) or neutral loss (highlight in green).

Based on previous reported crosstalk between O-GlcNAcylation and phosphorylation (36), we further compared the O-GlcNAc datasets with phospho dataset (20). Out of the 489 identified O-GlcNAc targets, a substantial 436 of them were also found to undergo phosphorylation (Fig.5B). On the other hand, seventeen proteins with kinase domains were identified as O-GlcNAc targets, so O-GlcNAcylation may affect the phosphorylation status by modulating the kinase activity. These observations underscore the intricate regulatory crosstalk.

Next, we explored whether the modifications occurred at the same sites by examining the data. Interestingly, we found certain proteins that were modified by both O-GlcNAcylation and O-fucosylation at precisely the same sites (supplementary data 6). For instance, a peptide from KINASE-RELATED PROTEIN OF UNKNOWN FUNCTION was found to be modified at the Serine 748 position by both O-GlcNAcylation and O-fucosylation (Fig 5C and 5D). To further map these sugar modifications, additional supporting data, particularly from ETD and EThcD analyses, would be valuable for in-depth investigation.

### Complexity with EThcD for site assignment

Despite significant efforts, assigning O-GlcNAc sites remains challenging due to two main factors. First, the modified peptide sequence, often found in intrinsic domains, tends to feature clusters of serine and threonine sites and requires significantly more fragments for unambiguous site assignment (15). Second, despite ETD, EtCID and EtHCD show promise for site assignment, their sensitivity is compromised by longer cycle times (25, 37). Furthurmore, it is noteworthy that in HCD activation, the glycosidic bond of O-GlcNAc is more labile to cleavage compared to that of N-GlcNAc.

To address these challenges, we employed EtHCD-m1, which considers both the loss and retention of the modification on fragments. The Protein Prospector software treats these options separately and reports higher scoring results. In cases where EtHCD-m1 scores are comparable, both possibilities are presented. If loss scores predominate, site information is excluded. To capture more site information, we also employed EThcD-m2, assuming all fragments are modified on S/T/N. As shown in Fig. 6A and 6B, when only assuming the fragments are modified, fragments (colored in blue and black) are assigned, but those with neutral loss (colored in green) remain unexplained. Fig.6A reveals a small number of unexplained fragments, whereas Fig.6B displays a more substantial quantity.

**Fig 6:**
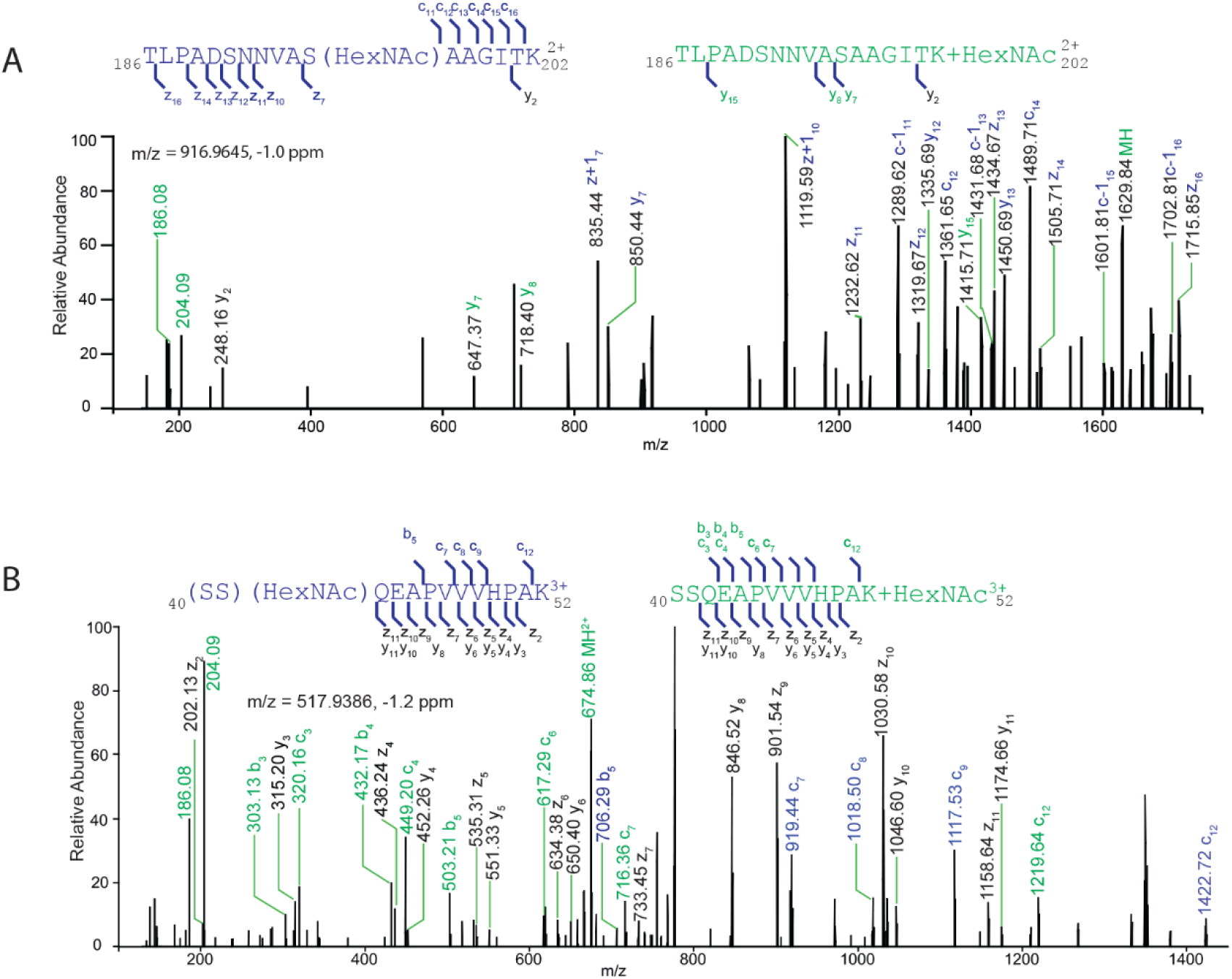
Complexity of site assignment using EtHCD. A. Fragments generated by ETD activation retain the modifications, whereas HCD fragments do not. The m/z 916.9645 identifies an O-GlcNAcylated peptide from ECT2 protein modified at Serine 196. B. While some fragments are detected modified owing to ETD, but many more fragments unmodified owing to HCD activation. The m/z 517.9380 identified an O-GlcNAc modified peptide from ECT5 protein modified at Serine 40 or 41. The fragments retaining modifications are highlighted in blue, while the fragments that undergo neutral loss are highlighted in green. The ions supporting both annotations are colored black.

These findings underscore the intricacies of site assignment (38), where observing a fragment modified due to ETD, but its equivalent remaining unmodified during HCD, can introduce ambiguity. To further enhance site assignment accuracy and best benefit from the rich spectrum generated by EThcD, future algorithm advancements and improved scoring methods are necessary. By addressing these challenges, we can continue to improve our understanding of O-GlcNAcylation and its functional implications in cellular processes.

## DISCUSSION

Recent discoveries have introduced several intriguing questions into our understanding of O-GlcNAcylation. Notably, the identification of atypical OGT in wheat (14), the mild phenotype observed in the *sec5* mutant (8), and the uncertain in vivo OGT activity of SPY, collectively raise inquiries regarding the prominence of SEC as a primary contributor to O-GlcNAcylation. Additionally, the substantial difference in expression levels between SEC (undetectable) and SPY(IBAQ=22.47, akin to AT1G60140 (trehalose phosphate synthase) or AT3G51470 (Protein phosphatase 2C) (39) and corroborated by our unpublished data, encourages an exploration of SEC’s functional significance.

We first confirmed that SEC and SPY have redundant functions during embryogenesis and that O-Glycosylation plays a crucial role in maintaining embryo cell viability. The failure to recover the *sec spy* double mutant despite extensive efforts underscores the functional overlap between these two genes.

SEC’s in vitro O-GlcNAcylation activity has been demonstrated, and transgenic DELLA proteins were found to exhibit reduced O-GlcNAc levels in *sec* mutant (11, 40). In our comprehensive quantitative analysis, we identified a significant reduction in O-GlcNAc levels within *sec-5* mutants. Utilizing both Data-Dependent Acquisition (DDA) and targeted quantification through Parallel Reaction Monitoring (PRM), we established that SEC predominantly drives O-GlcNAcylation in plants. Despite observing comparative lower O-GlcNAc enrichment in Arabidopsis seedlings compared to previous flower tissues, our SILIA labeling approach at seedling stage enabled precise quantification, confirming SEC’s major contribution to O-GlcNAcylation in vivo.

Detecting O-GlcNAc modification is challenged by its substoichiometry. For instance, in unenriched samples, only 126 O-GlcNAc modified peptides were detected among 14 million spectra (41). To expand the plant O-GlcNAc repertoire, we combined LWAC enrichment with extensive high pH reverse HPLC fractionations, which enabled the identification of 227 new targets, expanding our O-GlcNAcome database to encompass a total of 489 targets.

Alternative approaches using click chemistry have been employed to detect O-GlcNAc targets in Arabidopsis seedlings (18). By supplementing GalNAz and GlcNAz, metabolic glycan labeling was induced in Arabidopsis seedlings. Given GlcNAz’s toxicity, the enrichment focused on GalNAz-labeled samples. Subsequent click chemistry and enrichment through streptavidin beads enabled the profiling of 645 O-GlcNAc targets. Our comparative analysis revealed only 68 overlaps in seedling datasets and 75 in overall datasets. Notably, click chemistry enriches targets predominantly associated with cellular amide metabolic processes (18).

Given the observed redundant functions of O-GlcNAc and O-fucosylation during embryo stages, and the mild phenotype of SPY and SEC post-embryo development, the data strongly suggest an extensive crosstalk between O-GlcNAc and O-fucosylation. This suggests the possibility that these modifications may target the same substrates. Our comprehensive profiling of O-GlcNAc datasets has significantly expanded the number of overlapping substrates to 189, encompassing almost one-third of the detected targets. Remarkably, aside from their previously documented contrasting effects on the DELLA group (11, 40), recent findings illustrate that SEC and SPY collaboratively regulate style development through the bHLH transcription factor SPATULA (42). These new insights uncover novel roles for O-glycosylation and synergistic effects of these two sugar modifications.

EThcD demonstrates promising potential for site assignment of O-GlcNAc peptides and identifications. However, we also observed complexities in site assignment using EThcD due to the presence of both modified fragments from ETD and equivalent unmodified fragments resulting from HCD neutral loss. Presently, efficient identification of O-GlcNAc modified peptides, let alone site assignment, remains limited in available software. Addressing the intricacies of EThcD data, involving both modified and unmodified fragments, poses challenges and data ambiguity during searches. The enhancement of algorithms in the future holds substantial promise for refining modification localization and advancing research in this domain.

## DATA AVAILABILITY

The mass spectrometry proteomics data have been deposited to the ProteomeXchange Consortium via the PRIDE partner repository with the dataset identifier PXD044668, PXD044832, PXD044831, PXD044675, PXD044701. All other data are available from the corresponding author on reasonable request.

## Acknowledgments

We would like to thank Pei-qiao Xie for critical reading and editing of the manuscript. We would like to thank Robert Chalkley and Peter Baker for providing technical expertise and valuable comments. We would also like to thank Prof. Neil Olszewski for sharing the *spy* and *sec* mutants. This work was funded by the National Institutes of Health grants R01GM135706 and S10OD030441 to S.-L.X. and by the Carnegie Endowment Fund to the Carnegie Mass Spectrometry Facility. Author contributions – S.-L.,X. conceptualization; R.S., S.K. LWAC enrichment; R.S., S.K., T.G., A.V.R. data analysis; S.K. genetic crosses and analysis. R.S., T.G., A.V.R. Figures. S-L,X manuscript writing.

## Experimental Procedures

### Plant growth and tissue

Wild-type (Col), *sec-5* (SALK_034290) and *spy-3*, and *sec-5* (-/-) *spy-3*(+/-) were grown in the greenhouse for 30-40 days. Siliques were taken and examined under the Amscope MD500 microscope. Genotyping primers (*sec-5* genotyping primers-JMLB1-GGCAATCAGCTGTTGCCCGTCTCACTGGTG, SEC RF1-tcccgacctgtctttctttccgat, SEC RR1-ttccacgcgtttgccatgtttc, *spy-3* genotyping primers-JP91-GCGACCTATCACCATTGGA, SPY LPD3-TCGACCTGCCTGCAATCAAA).

### SILIA Labeling

The wild type (Col) and *sec* seedlings were grown on Hoagland medium containing ^14^N or ^15^N isotope salt (1.34 g/L Hoagland’s No. 2 salt mixture without nitrogen (Caisson labs), 6 g/L Phytoblend, and 1 g/L KNO_3_ or 1 g/L K^15^NO3 (Cambridge Isotope Laboratories), pH 5.8) for 14 days. Plates were placed vertically in a growth chamber under constant light conditions at 22 °C. Similarly, a single biological replicate of Col and *spy-4* were labeled by ^14^N and ^15^N isotopes, respectively, for subsequent large-scale identification of O-GlcNAc peptide.

### SILIA sample Preparation

Whole plant tissues were harvested and cryomilled in liquid nitrogen. For quantifications, the tissues from six samples were equally mixed to make 3 biological replicates for both wild-type and *sec-5* mutant (F1: ^14^N Col /^15^N *sec-5*; F2:^14^N Col /^15^N *sec-5*; R1, ^15^N Col/ ^14^N *sec-5*). Similarly, one replicate was prepared for wild-type and *spy-4* mutants (S1: ^14^N Col /^15^N *spy-4)*. Protein extraction was performed using phenol extraction method as described by (20). Following protein extraction, proteins were reduced and alkylated followed by trypsin digestion. Peptides were desalted using Sep-PAK C18 cartridges (Waters).

### LWAC enrichment and high pH reverse fractionation

The LWAC column was packed as described (20), with slight modification for three-round glycosylated peptide enrichment. Peptides were resuspended in 100 µl LWAC buffer (100 mM Tris, pH 7.5, 150 mM NaCl, 2 mM MgCl_2_, 2 mM CaCl_2_, 5% (vol/vol) acetonitrile). Chromatography was performed at a flow rate of 100 µl/min, with a single glycopeptide-enriched tail collected after 1.3 ml.

To prevent column overload, in the initial LWAC enrichment, total peptides were divided into aliquots of ∼1-2 mg each and run separately. In Col/*sec-5* experiments, the starting peptides were as follows: F1: 11 mg (split into 6 runs with 1.8 mg injection each), F2: 5 mg (split into 5 runs with 1 mg injection each), R1: 5 mg (split into 5 runs with 1 mg injection each). For S1 Col/*spy-4* experiment, the starting peptides were 20 mg (split into 10 runs with 2 mg injection each).The glycopeptide-enriched fractions were combined and desalted for the subsequent round. Subsequent rounds utilized peptides from the prior round, with the same glycopeptide-enriched fraction being targeted. Column cleaning involved injecting 100 μl of 40 mm GlcNAc in the LWAC buffer to elute any bound glycopeptides.

The high pH reverse HPLC fractionations were performed as described (20) and collected as described in supplementary data 8. For F1: ^14^N Col /^15^N *sec-5*, 8 fractions were collected. For F2:^14^N Col /^15^N *sec-5*, 18 fractions were collected for DDA assay, and 10 of these underwent targeted assay. For R1:, ^15^N Col/ ^14^N *sec-5*, 10 fractions collected and subjected to targeted assay. For ^14^N Col/^15^N *spy-4*, 52 fractions were collected and analyzed through DDA analysis.

### Nanoflow LC-MS/MS

Peptides were analyzed by liquid chromatography–tandem mass spectrometry (LC-MS/MS) on an Easy LC 1200 UPLC liquid chromatography system connected to Q-Exactive HF hybrid quadrupole-Orbitrap mass spectrometer or Orbitrap Eclipse with ETD (Thermo Fisher) (Supplemental Data 7). Peptides were separated using Easy-Spray C18 columns (75 μm × 50 cm) (Thermo, ES803A or ES903) or Ion opticks column Aurora UHPLC Emitter column with nanoZero (AUR2-25075C18A), with the exception for F2/R2 datasets, which utilized a trapping column before separation (NanoViper trap column Acclaim pepMap C18 column (75 μm × 2 cm, ThermoFisher, 164946)). The flow rate was 300 nL/min, and a 120 min gradient was employed. Peptides were eluted by a gradient from 3 to 28% solvent B (80% (v/v) acetonitrile/0.1% (v/v) formic acid) over 100 min, followed by an increase to 44% solvent B over 20 min, and concluded with by a short wash at 90% solvent B.

For data acquired on Q-Exactive HF with DDA acquisition, precursor scan was from mass-to-charge ratio (m/z) 375 to 1600 (resolution 120,000; AGC 3.0E6, maximum injection time 100 ms) and top 20 most intense multiply charged precursors were selected for fragmentation (resolution 15,000, AGC 5E4, maximum injection time 60 ms, isolation window 1.0 m/z, scan range 200-2000, spectrum data type profile, minimum AGC target 1.2e3, intensity threshold 2.0 e4, include charge state =2-8) Peptides were fragmented with higher-energy collision dissociation (HCD) with normalized collision energy (NCE) 27.

For data acquired on Q-Exactive HF with PRM acquisition, data were acquired in the PRM mode. Peptides were similarly separated as described above. The samples were analyzed using the PRM mode with an isolation window of 1.4 Th. PRM scans were done using 60,000 resolution (AGC target 2e5, 200ms maximum injection time) triggered by an inclusion list. Normalized collision energy 27 was used in HCD mode. The PRM data were analyzed using the skyline software (22, 43).

For HCD/EThcD data acquired on an Orbitrap Eclipse (Thermo Scientific, San Jose, CA, USA), the mass spectrometer was coupled with an Easy LC 1200 UPLC liquid chromatography system (Thermo Fisher). Peptides were separated using LC gradient the same described as above. The precursor ions were scanned from 375 to 1600 m/z (resolution 120,000; AGC 4.0E5) and the charge state 2^+^ to 6^+^ were filtered in the quadrupole with a selection window of 1.0 m/z and MIPS Peptide filter enabled. Peptides were fragmented with higher energy collision dissociation (HCD) with normalized collision energy (NCE) 27. Dynamic exclusion was enabled for 10s. For Triggered EtHCD, mass trigger was set as 204.0866 with mass tolerance 25ppm. Ions were isolated in quadrupole, EThcd with activated (resolution 30,000, AGC target 5E4, Normalized AGC target 100%, microscans=3, with supplemental collision energy 35).

### Data Analysis (SILIA)

MS/MS data was processed using PAVA script, which generates centroid MS2 peaklist. The data analysis was analyzed as described in (21) but reports the quantification in peptide level. For data acquired on Eclipse, the peaklists were split into HCD and EThcd peaklist categories. Data were searched using Protein Prospector against the TAIR database Arabidopsis thaliana from December 2010 (https://www.arabidopsis.org/), concatenated with sequence randomized versions of each protein (a total of 35386 entries). A precursor mass tolerance was set to 10 ppm and MS/MS2 tolerance was set to 20 ppm., Carbamidomethylcysteine was specified as a constant modification, while the variable modifications include protein N-terminal acetylation, peptide N-terminal Gln conversion to pyroglutamate, Met oxidation. Additionally, GlcNAc modifications were included: either HexNAc modification of serine and threonine, asparagine and neutral loss of HexNAc (method named as HCD-m or EtHCD-m1; HCD-m considers b, y ions while EtHCD-m1 takes into account c, z, b, y ions.) or HexNAc modification of serine, threonine and asparagine (method named as EtHCD-m2 which considers c, z, b, y ions). The false discovery rate (FDR) was set at 1% for both proteins and peptides. 204 filtered lists from HCD peaklists were used for search, alongside the complete peaklists. The cleavage specificity was set to trypsin, allowing up to two missed cleavages and a maximum of three modifications. PRM data quantification was described as in (22).

### Bioinformatic analysis

Subcellular localization analysis was performed with SUBA5.0 (https://suba.live/) with default settings (31). The consensus subcellular localization was used. GO analysis was performed using the “Database for Annotation, Visualization and Integrated Discovery” (DAVID) application (DAVID Functional Annotation Bioinformatics Microarray Analysis) (44) with the Arabidopsis thaliana fully annotated protein library as the background.

## Supplemental information

Supplemental Data 1: SILIA quantification of O-GlcNAc levels in F1 and F2 replicates, comparing wild-type (^14^N) vs *sec-5* mutant (^15^N).

Supplemental Data 2: SILIA quantification of O-GlcNAc levels in F2 and R1 recipical -labeled replicates comparing wild-type vs *sec-5* mutant.

Supplemental Data 3: List of detected O-GlcNAc peptides.

Supplemental Data 4: List of identified O-GlcNAc proteins, compared to other O-GlcNAc and O-fucosylation datasets.

Supplemental Data 5: SUBA analysis of O-GlcNAc targets.

Supplemental Data 6: Common modification sites detected between O-GlcNAc and O- fucoslation.

Supplemental Data 7: LC-MS method, search, and quantification details for different experiments.

Supplemental Data 8: Summary of Data Files.

---

**Supplemental Fig S1:**
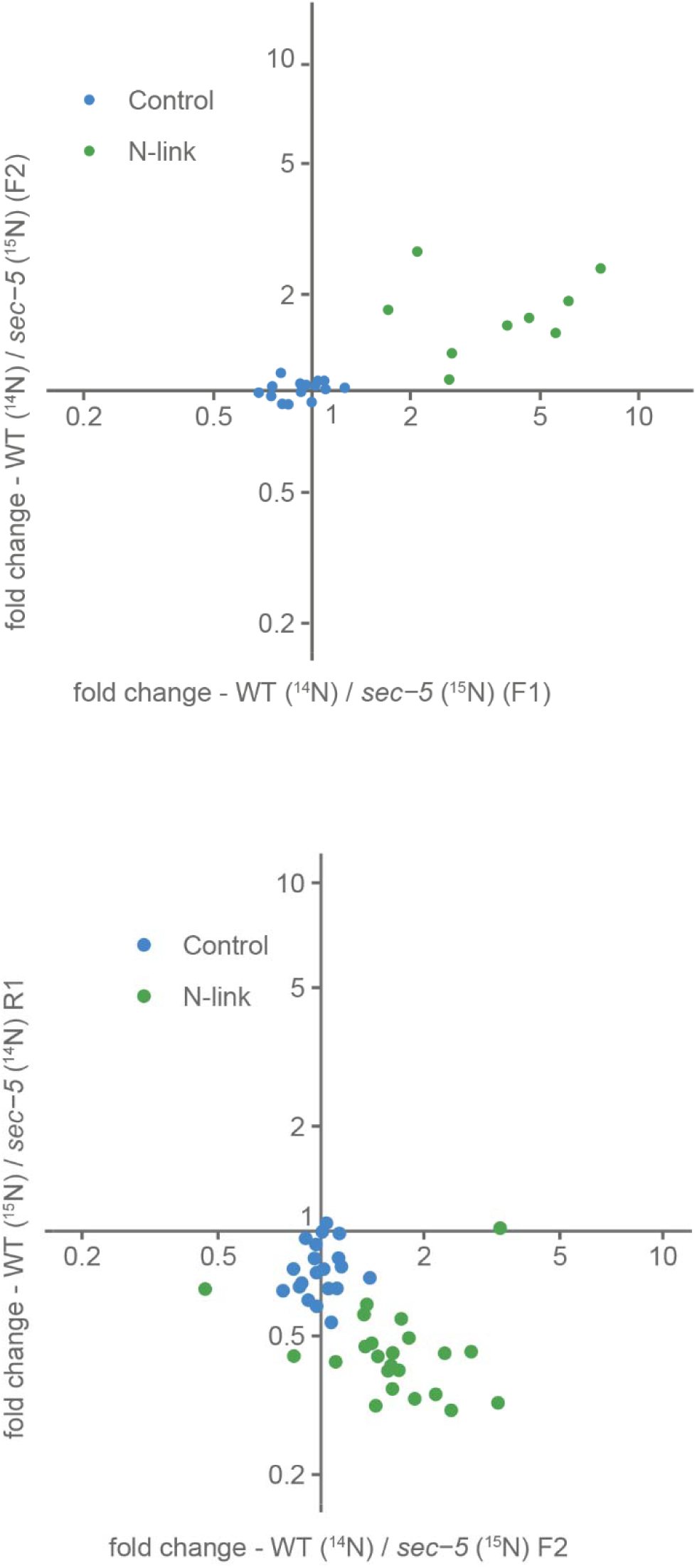
N-linked peptides quantifications show no significant differences between wild-type and *sec-5* mutant. The top panel displays MS1 quantification from the F1 and F2 datasets. The bottom panel shows PRM quantification from the F2 and R1 datasets. Control peptides are colored blue, while N-linked peptides are colored green.

**Supplemental Fig S2:**
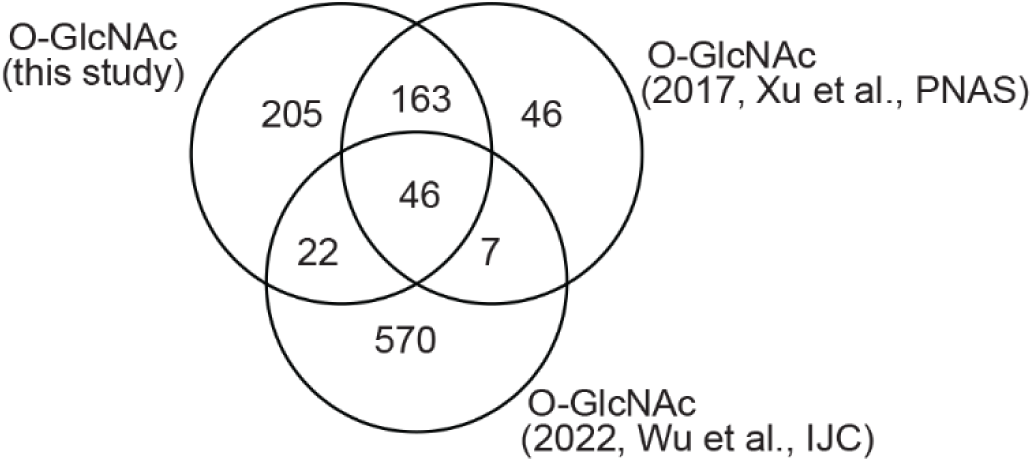
Overlap of our O-GlcNAc sets using seedlings vs. our previous O-GlcNAc datasets from inflorescence tissues, vs. O-GlcNAc datasets from seedlings using click chemistry from Wu et al.

## Notes

### Competing Interest Statement

The authors have declared no competing interest.

### Summary of Updates

The authorship list order was incorrect (the PDF version was correct). I, the corresponding author, should be last instead of first. Sorry for the mistake.

## REFERENCES

1. Ma, J., Wu, C., and Hart, G. W. (2021) Analytical and Biochemical Perspectives of Protein O-GlcNAcylation. Chem. Rev. 121, 1513–1581

2. Gambetta, M. C., Oktaba, K., and Müller, J. (2009) Essential role of the glycosyltransferase sxc/Ogt in polycomb repression. Science 325, 93–96

3. Bond, M. R., and Hanover, J. A. (2013) O-GlcNAc cycling: a link between metabolism and chronic disease. Annu. Rev. Nutr. 33, 205–229

4. Lagerlöf, O., Slocomb, J. E., Hong, I., Aponte, Y., Blackshaw, S., Hart, G. W., and Huganir, R. L. (2016) The nutrient sensor OGT in PVN neurons regulates feeding. Science 351, 1293–1296

5. Biwi, J., Biot, C., Guerardel, Y., Vercoutter-Edouart, A.-S., and Lefebvre, T. (2018) The Many Ways by Which O-GlcNAcylation May Orchestrate the Diversity of Complex Glycosylations. Molecules 23

6. Jacobsen, S. E., and Olszewski, N. E. (1993) Mutations at the SPINDLY locus of Arabidopsis alter gibberellin signal transduction. Plant Cell 5, 887–896

7. Chen, D., Juárez, S., Hartweck, L., Alamillo, J. M., Simón-Mateo, C., Pérez, J. J., Fernández-Fernández, M. R., Olszewski, N. E., and García, J. A. (2005) Identification of secret agent as the O-GlcNAc transferase that participates in Plum pox virus infection. J. Virol. 79, 9381–9387

8. Xing, L., Liu, Y., Xu, S., Xiao, J., Wang, B., Deng, H., Lu, Z., Xu, Y., and Chong, K. (2018) O-GlcNAc transferase SEC activates histone methyltransferase ATX1 to regulate flowering. EMBO J. 37

9. Hartweck, L. M., Genger, R. K., Grey, W. M., and Olszewski, N. E. (2006) SECRET AGENT and SPINDLY have overlapping roles in the development of Arabidopsis thaliana L. Heyn. J. Exp. Bot. 57, 865–875

10. Hartweck, L. M., Scott, C. L., and Olszewski, N. E. (2002) Two O-linked N-acetylglucosamine transferase genes of Arabidopsis thaliana L. Heynh. have overlapping functions necessary for gamete and seed development. Genetics 161, 1279–1291

11. Zentella, R., Sui, N., Barnhill, B., Hsieh, W.-P., Hu, J., Shabanowitz, J., Boyce, M., Olszewski, N. E., Zhou, P., Hunt, D. F., and Sun, T.-P. (2017) The Arabidopsis O-fucosyltransferase SPINDLY activates nuclear growth repressor DELLA. Nat. Chem. Biol. 13, 479–485

12. Bi, Y., Shrestha, R., Zhang, Z., Hsu, C.-C., Reyes, A. V., Karunadasa, S., Baker, P. R., Maynard, J. C., Liu, Y., Hakimi, A., Lopez-Ferrer, D., Hassan, T., Chalkley, R. J., Xu, S.-L., and Wang, Z.-Y. (2023) SPINDLY mediates O-fucosylation of hundreds of proteins and sugar-dependent growth in Arabidopsis. Plant Cell 35, 1318–1333

13. Zentella, R., Wang, Y., Zahn, E., Hu, J., Jiang, L., Shabanowitz, J., Hunt, D. F., and Sun, T.-P. (2023) SPINDLY O-fucosylates nuclear and cytoplasmic proteins involved in diverse cellular processes in plants. Plant Physiol. 191, 1546–1560

14. Fan, M., Miao, F., Jia, H., Li, G., Powers, C., Nagarajan, R., Alderman, P. D., Carver, B. F., Ma, Z., and Yan, L. (2021) O-linked N-acetylglucosamine transferase is involved in fine regulation of flowering time in winter wheat. Nat. Commun. 12, 2303

15. Ma, J., Hou, C., and Wu, C. (2022) Demystifying the O-GlcNAc Code: A Systems View. Chem. Rev. 122, 15822–15864

16. Wulff-Fuentes, E., Berendt, R. R., Massman, L., Danner, L., Malard, F., Vora, J., Kahsay, R., and Olivier-Van Stichelen, S. (2021) The human O-GlcNAcome database and meta-analysis. Sci Data 8, 25

17. Li, X., Lei, C., Song, Q., Bai, L., Cheng, B., Qin, K., Li, X., Ma, B., Wang, B., Zhou, W., Chen, X., and Li, J. (2023) Chemoproteomic profiling of O-GlcNAcylated proteins and identification of O-GlcNAc transferases in rice. Plant Biotechnol. J. 21, 742–753

18. Wu, J., Lei, C., Li, X., Dong, X., Qin, K., Hong, W., Li, J., Zhu, Y., and Chen, X. (2023) Chemoproteomic profiling of O-GlcNAcylation in Arabidopsis Thaliana by using metabolic glycan labeling. Isr. J. Chem. 63

19. Xu, S., Xiao, J., Yin, F., Guo, X., Xing, L., Xu, Y., and Chong, K. (2019) The Protein Modifications of -GlcNAcylation and Phosphorylation Mediate Vernalization Response for Flowering in Winter Wheat. Plant Physiol. 180, 1436–1449

20. Xu, S.-L., Chalkley, R. J., Maynard, J. C., Wang, W., Ni, W., Jiang, X., Shin, K., Cheng, L., Savage, D., Hühmer, A. F. R., Burlingame, A. L., and Wang, Z.-Y. (2017) Proteomic analysis reveals O-GlcNAc modification on proteins with key regulatory functions in Arabidopsis. Proc. Natl. Acad. Sci. U. S. A. 114, E1536–E1543

21. Shrestha, R., Reyes, A. V., Baker, P. R., Wang, Z.-Y., Chalkley, R. J., and Xu, S.-L. (2022) 15N Metabolic Labeling Quantification Workflow in Arabidopsis Using Protein Prospector. Front. Plant Sci. 13, 832562

22. Reyes, A. V., Shrestha, R., Baker, P. R., Chalkley, R. J., and Xu, S.-L. (2022) Application of Parallel Reaction Monitoring in 15N Labeled Samples for Quantification. Front. Plant Sci. 13, 832585

23. Holstein Sherwood, C. A., Gafken, P. R., and Martin, D. B. (2011) Collision energy optimization of b- and y-ions for multiple reaction monitoring mass spectrometry. J. Proteome Res. 10, 231–240

24. Pino, L. K., Searle, B. C., Bollinger, J. G., Nunn, B., MacLean, B., and MacCoss, M. J. (2020) The Skyline ecosystem: Informatics for quantitative mass spectrometry proteomics. Mass Spectrom. Rev. 39, 229–244

25. Yu, Q., Wang, B., Chen, Z., Urabe, G., Glover, M. S., Shi, X., Guo, L.-W., Kent, K. C., and Li, L. (2017) Electron-Transfer/Higher-Energy Collision Dissociation (EThcD)-Enabled Intact Glycopeptide/Glycoproteome Characterization. J. Am. Soc. Mass Spectrom. 28, 1751–1764

26. Riley, N. M., and Coon, J. J. (2018) The Role of Electron Transfer Dissociation in Modern Proteomics. Anal. Chem. 90, 40–64

27. Frese, C. K., Altelaar, A. F. M., van den Toorn, H., Nolting, D., Griep-Raming, J., Heck, A. J. R., and Mohammed, S. (2012) Toward full peptide sequence coverage by dual fragmentation combining electron-transfer and higher-energy collision dissociation tandem mass spectrometry. Anal. Chem. 84, 9668–9673

28. Trinidad, J. C., Barkan, D. T., Gulledge, B. F., Thalhammer, A., Sali, A., Schoepfer, R., and Burlingame, A. L. (2012) Global identification and characterization of both O-GlcNAcylation and phosphorylation at the murine synapse. Mol. Cell. Proteomics 11, 215–229

29. Janetzko, J., and Walker, S. (2014) The making of a sweet modification: structure and function of O-GlcNAc transferase. J. Biol. Chem. 289, 34424–34432

30. Stephen, H. M., Adams, T. M., and Wells, L. (2021) Regulating the Regulators: Mechanisms of Substrate Selection of the O-GlcNAc Cycling Enzymes OGT and OGA. Glycobiology 31, 724–733

31. Hooper, C., Millar, H., Black, K., Castleden, I., Aryamanesh, N., and Grasso, S. (2022) Subcellular Localisation database for Arabidopsis proteins version 5. The University of Western Australia

32. Molitor, L., Bacher, S., Burczyk, S., and Niessing, D. (2021) The Molecular Function of PURA and Its Implications in Neurological Diseases. Front. Genet. 12, 638217

33. Bi, Y., Deng, Z., Ni, W., Shrestha, R., Savage, D., Hartwig, T., Patil, S., Hong, S. H., Zhang, Z., Oses-Prieto, J. A., Li, K. H., Quail, P. H., Burlingame, A. L., Xu, S.-L., and Wang, Z.-Y. (2021) Arabidopsis ACINUS is O-glycosylated and regulates transcription and alternative splicing of regulators of reproductive transitions. Nat. Commun. 12, 945

34. Tan, Z.-W., Fei, G., Paulo, J. A., Bellaousov, S., Martin, S. E. S., Duveau, D. Y., Thomas, C. J., Gygi, S. P., Boutz, P. L., and Walker, S. (2020) O-GlcNAc regulates gene expression by controlling detained intron splicing. Nucleic Acids Res. 48, 5656–5669

35. Mutanwad, K. V., and Lucyshyn, D. (2022) Balancing O-GlcNAc and O-fucose in plants. FEBS J. 289, 3086–3092

36. Hart, G. W., Slawson, C., Ramirez-Correa, G., and Lagerlof, O. (2011) Cross talk between O-GlcNAcylation and phosphorylation: roles in signaling, transcription, and chronic disease. Annu. Rev. Biochem. 80, 825–858

37. Frese, C. K., Zhou, H., Taus, T., Altelaar, A. F. M., Mechtler, K., Heck, A. J. R., and Mohammed, S. (2013) Unambiguous phosphosite localization using electron-transfer/higher-energy collision dissociation (EThcD). J. Proteome Res. 12, 1520–1525

38. Maynard, J. C., and Chalkley, R. J. (2021) Methods for Enrichment and Assignment of N-Acetylglucosamine Modification Sites. Mol Cell Proteomics 20, 100031

39. Mergner, J., Frejno, M., List, M., Papacek, M., Chen, X., Chaudhary, A., Samaras, P., Richter, S., Shikata, H., Messerer, M., Lang, D., Altmann, S., Cyprys, P., Zolg, D. P., Mathieson, T., Bantscheff, M., Hazarika, R. R., Schmidt, T., Dawid, C., Dunkel, A., Hofmann, T., Sprunck, S., Falter-Braun, P., Johannes, F., Mayer, K. F. X., Jürgens, G., Wilhelm, M., Baumbach, J., Grill, E., Schneitz, K., Schwechheimer, C., and Kuster, B. (2020) Mass-spectrometry-based draft of the Arabidopsis proteome. Nature 579, 409–414

40. Zentella, R., Hu, J., Hsieh, W.-P., Matsumoto, P. A., Dawdy, A., Barnhill, B., Oldenhof, H., Hartweck, L. M., Maitra, S., Thomas, S. G., Cockrell, S., Boyce, M., Shabanowitz, J., Hunt, D. F., Olszewski, N. E., and Sun, T.-P. (2016) O-GlcNAcylation of master growth repressor DELLA by SECRET AGENT modulates multiple signaling pathways in Arabidopsis. Genes Dev. 30, 164–176

41. Hahne, H., Sobotzki, N., Nyberg, T., Helm, D., Borodkin, V. S., van Aalten, D. M. F., Agnew, B., and Kuster, B. (2013) Proteome wide purification and identification of O-GlcNAc-modified proteins using click chemistry and mass spectrometry. J. Proteome Res. 12, 927–936

42. Jiang, Y., Curran-French, S., Koh, S. W. H., Jamil, I., Argirò, L., Lopez, S., Martins, C., Saalbach, G., and Moubayidin, L. (2023) Post-translational modification of SPATULA by SECRET AGENT and SPINDLY promotes organ symmetry transition at the gynoecium apex. bioRxiv

43. MacLean, B., Tomazela, D. M., Shulman, N., Chambers, M., Finney, G. L., Frewen, B., Kern, R., Tabb, D. L., Liebler, D. C., and MacCoss, M. J. (2010) Skyline: an open source document editor for creating and analyzing targeted proteomics experiments. Bioinformatics 26, 966–968

44. Huang, D. W., Sherman, B. T., Tan, Q., Collins, J. R., Alvord, W. G., Roayaei, J., Stephens, R., Baseler, M. W., Lane, H. C., and Lempicki, R. A. (2007) The DAVID Gene Functional Classification Tool: a novel biological module-centric algorithm to functionally analyze large gene lists. Genome Biol 8, R183

